# Regulation of *Chlamydia trachomatis* infection in the female genital tract by type I and type II interferons

**DOI:** 10.64898/2026.07.17.739123

**Authors:** Rongze He, Yi Wu, Ahmed Abdelsalam, Yihui Wang, Huizhou Fan, Guangming Zhong

**Author notes:** These authors contributed equally to the work reported in this manuscript (Rongze He, Yi Wu). Corresponding author: Guangming Zhong, Department of Microbiology, Immunology and Molecular Genetics, University of Texas Health Science Center at San Antonio, 7703 Floyd Curl Drive, San Antonio, Texas 78229, USA.

## Abstract

Following an intravaginal inoculation with *Chlamydia trachomatis*, mice deficient in type I interferon receptor IFNαR1 (IFNαR1^-/-^) significantly increased the yield of live chlamydiae on days 3 & 5 but reduced it to the level of wild-type mice by day 7, while mice deficient in type II interferon receptor IFNγR1 (IFNγR1^-/-^) significantly increased the chlamydial yield by day 5 and the increase persisted throughout the remainder of the infection course. These observations reveal a temporal division of labor between type I & II interferons in regulating *C. trachomatis* infection in the female genital tract. Interestingly, mice deficient in both IFNαR1 & IFNγR1 exhibited higher mortality and shed more chlamydial organisms than IFNγR1^-/-^ mice by week 6, suggesting that IFNαR1 remains critical for inhibiting *C. trachomatis* at late stages. An anti-IFNαR1 antibody blockade significantly increased chlamydial yields in IFNγR1^-/-^ mice, suggesting that the anti-chlamydial activities of type I & II interferon systems are both distinct and overlapping throughout the infection course. Furthermore, the anti-chlamydial activity of type I interferon signaling is localized to the lower vagina, while that of type II interferon signaling is localized to the upper vagina. Thus, we have demonstrated that type I & II interferons function complementarily and synergistically in time and space to control *C. trachomatis* infection, laying the foundation for further elucidating the mechanisms of IFN regulation of chlamydial infection and for developing interventional and preventive strategies against *C. trachomatis* in the female genital tract.

**Importance:** Lack of information on the precise roles of type I and type II interferons during chlamydial infection has hindered the development of interferon-based strategies to prevent chlamydial infection and pathogenicity. The current study has revealed a temporal division of labor between type I & type II interferons in regulating chlamydial infection in the female genital tract, with type I acting earlier than type II during the innate phase. Nevertheless, type I remains critical for cooperating with type II to suppress chlamydia 6 weeks after infection. Finally, type I interferons seem to mainly target chlamydial infection in the lower vagina, while type II interferons target the infection in the upper vagina. These new findings on the distinct and overlapping roles of type I and type II interferons in regulating chlamydial infection may guide the development of interferon-based precision strategies to reduce chlamydial infection and pathogenicity in the female genital tract.

## Introduction

Sexually transmitted *Chlamydia trachomatis* infection in the lower genital tract may lead to upper genital tract complications such as pelvic inflammatory disease and tubal infertility (1, 2). The precise mechanisms of mucosal immunity for controlling *C. trachomatis* infection in the female genital tract remain unclear. The genital mucosal immunity may consist of the following three layers (3): Pre-existing effectors that are expressed constitutively (4) and can act on microbes within the 4 hours of infection, the early-induced responses that are activated by pattern recognition receptor (PRR) detection of microbial signals and amplified by cytokine receptors, which dominate the anti-infection activities within the first 4 days of infection (5), and finally, the adaptive immune responses that are mediated by microbial epitope-specific lymphocytes most likely induced in the draining lymph nodes, which may require 4 or more days to develop (6, 7).

Regardless of the stage of microbial infection, both type I (8) and type II (9, 10) interferons play critical roles in mucosal immunity (11, 12). *Chlamydia* has been shown to activate the DNA sensor signaling pathway consisting of cGAS (cyclic GMP–AMP synthase) and STING (stimulator of interferon genes) (13–17), leading to the production of type I interferons (17–19). Mice deficient in either cGAS or STING displayed significantly increased susceptibility to *C. trachomatis* infection in the female genital tract on days 3 and 5 post-infection (20). The increased susceptibility was recently reproduced in mice lacking IFNαR1 (IFNαR1^-/-^) (21), demonstrating a critical role for type I IFN signaling in controlling *C. trachomatis* infection in the female genital tract before adaptive immune responses are developed. However, the mechanism of IFNαR1-mediated anti-chlamydial activity remains unknown, and it is unclear whether IFNαR1 signaling still exerts anti-chlamydial activity at late stages of infection.

Type II interferon or IFNγ is the most effective cytokine for inhibiting intracellular replication of chlamydia in cultured cells (22) and in mice (23, 24), via multiple mechanisms (25–28). Mice deficient in either IFNγ or IFNγ receptor were highly susceptible to chlamydial infection (29). In addition to its cell-autonomous mechanisms (26), IFNγ may also exert anti-chlamydial activity indirectly, for example, by promoting antibody-mediated immunity (30). The cellular sources of IFNγ in the female genital tract are diverse (10). During the innate immune phase, NK cells are considered a significant source, as indicated by observations following anti-NK1.1 depletion (31). However, it was found that ex-ILC3s that also express NK1.1 may be a major contributor of innate IFNγ in the female genital tract (32) in addition to the contribution of ILC1s (33). During the adaptive immunity phase, chlamydia epitope-specific CD4^+^ T cells are considered the primary source of IFNγ in the female genital tract (29) or the small intestine (34). However, mice deficient in IFNγ only in CD4^+^ T cells showed genital shedding courses similar to those of wild-type control mice, suggesting that there are additional cellular sources of IFNγ during adaptive immunity (35, 36). Regardless of the precise cellular sources, IFNγ is, to date, the most reproducible correlate of protection in women exposed to *C. trachomatis* (37–39). Thus, it is imperative to further characterize IFNγ‘s anti-chlamydial activity in the female genital tract.

Since both type I and II interferons are induced by chlamydial infection and play prominent roles in controlling it, the current study aims to investigate how they interact with one another during chlamydial infection and across different genital tissue segments. We found that IFNαR1^-/-^ mice significantly increased the yield of live chlamydia on days 3 & 5 but reduced it to the level of wild-type mice by day 7, while IFNγR1^-/-^ mice significantly increased the chlamydial yield only by day 5, and the increase lasted for the remainder of the infection course. These results demonstrate a temporal division of labor between type I & II interferon-mediated anti-chlamydial activities in the female genital tract following an intravaginal infection with *C. trachomatis*. However, mice deficient in both IFNαR1 & IFNγR1 developed a higher mortality rate than IFNγR1^-/-^ mice and shed high levels of live chlamydial organisms throughout the infection course, whereas IFNγR1^-/-^ mice reduced their shedding 6 weeks post-infection. These observations suggest that IFNαR1 signaling may still play a critical role in inhibiting *C. trachomatis* at late stages of infection. This conclusion is supported by the observation that blocking IFNαR1 with an anti-IFNαR1 antibody in IFNγR1^-/-^ mice at week 6 post-infection significantly increased chlamydial yield. Thus, the anti-chlamydial activities of type I & II interferons are both distinct and overlapping throughout the course of infection. Furthermore, the anti-chlamydial activity of type I interferon signaling is mainly localized to the lower vaginal tissue, whereas that of type II interferon signaling is mapped to the upper vaginal and ectocervical tissues. These observations suggest that there is also a spatial division of labor between the anti-chlamydial activities of type I and II interferons. Together, our results have laid a foundation for further elucidation of how type I and II IFN systems regulate sexually transmitted infection by an obligate intracellular bacterium, and inform the development of therapeutic and preventive strategies against *C. trachomatis* infection in the female genital tract.

## Results

### 1. The early onset of type I IFNαR regulation of *Chlamydia trachomatis* infection is eventually taken over by type II IFNγR in the female genital tract

Chlamydial infection induces both type I & II interferons (10, 17–20, 30, 31, 40), and the effects of each on chlamydial infection in the female genital tract have been evaluated separately (21, 34, 41–48). The current study aimed to assess the potential co-operation of the two interferon types in regulating *C. trachomatis* infection in the genital tract. We first compared the effects of type I & II interferon signaling on chlamydial infection under three different inoculum doses (Fig. 1). Following an intravaginal inoculation with 2 x 10^5^ IFUs of *C. trachomatis*, wild-type (C57BL/6J) mice failed to shed any significant level of live *C. trachomatis* in the vaginal swabs. In contrast, substantial levels of live *C. trachomatis* were recovered from IFNαR1^-/-^ mice on days 3 & 5 (compared with those from wild-type mice), indicating a critical role of IFNαR1 signaling in inhibiting *C. trachomatis* infection. However, IFNγR1^-/-^ mice did not significantly increase the infectious yields of *C. trachomatis* in vaginal swabs on either day 3 or 5, indicating that IFNγR1 signaling is not as crucial as IFNαR1 signaling in controlling *C. trachomatis* infection during the early stages of infection. Mice deficient in both IFNαR1 and IFNγR1 signaling (double-knockout mice) showed a trend toward higher yields of live *C. trachomatis* but did not differ significantly from wild-type mice, likely due to substantial variation and the limited sample size. When the inoculum dose was elevated to 5 × 10^6^ IFUs, the wild-type mice were significantly infected with a peak shedding of live chlamydial organisms on day 3 postinfection. Importantly, IFNαR1^-/-^ mice further significantly increased the yield of live *C. trachomatis* on days 3 & 5, whereas IFNγR1^-/-^ mice significantly increased the yield only as early as day 5. These observations have validated the conclusion that the anti-chlamydial activity of IFNαR1 signaling occurs earlier than that of IFNγR1 signaling. Further, the yield of live *C. trachomatis* from the IFNαR1^-/-^ mice declined to the level observed in wild-type mice by day 7, whereas IFNγR1^-/-^ mice continued to increase significantly throughout the course of infection. These observations suggest that the early anti-chlamydial activity of IFNαR1 signaling is transient, whereas the anti-chlamydial activity of IFNγR1 signaling is sustained. This conclusion is further supported by the observation made at an inoculum dose of 2 × 10^8^ IFUs that IFNγR1^-/-^ mice maintained significantly high yields of live *C. trachomatis* for >6 weeks, while IFNαR1^-/-^ mice were no longer able to increase the yields of live *C. trachomatis* at any time point, probably due to the high inoculum dose-activated anti-chlamydial mechanisms that are independent of IFNαR1 signaling. Thus, the inhibition of *C. trachomatis* within the first 3 days after intravaginal infection primarily depends on IFNαR1 signaling at low and middle inoculation doses, but the early inhibitory activity is transient. The IFNαR1 signaling-dependent early anti-chlamydial activity is subsequently replaced by the IFNγR1 signaling-dependent mechanisms that persist for the remainder of the infection. The above observations collectively demonstrate a clear temporal division of labor between IFNαR1 and IFNγR1 signaling in controlling *C. trachomatis* infection in the female genital tract.

**Fig. 1.**
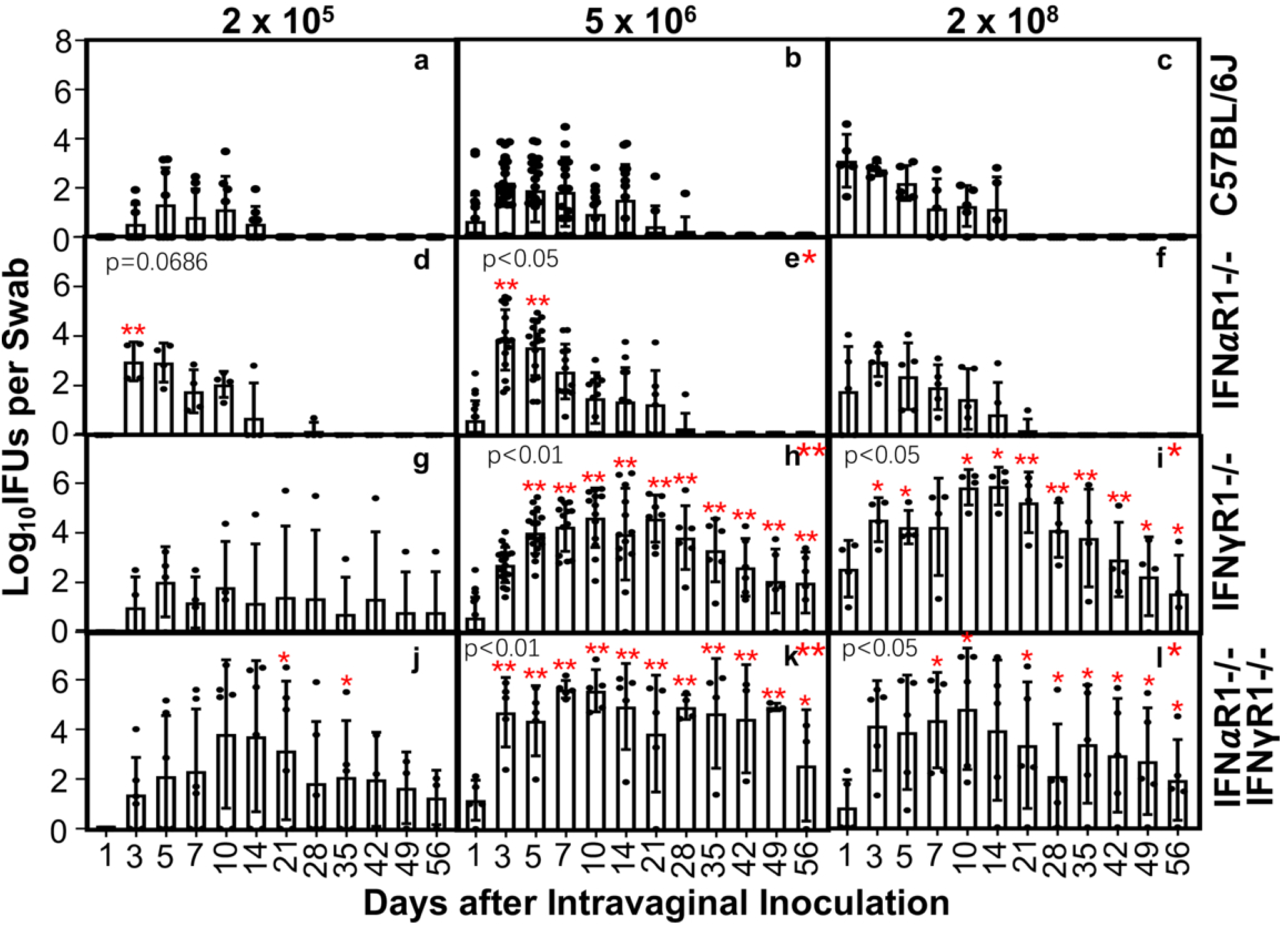
The effects of IFNαR and/or IFNγR signaling on *Chlamydia trachomatis* infection in the female mouse genital tract under various inoculation doses. Female mice without (C57BL/6J, n=5 to 22, panels a-c) or with deficiency in type I interferon receptor or interferon alpha/beta receptor 1 (IFNαR1^-/-^, n=4 to 17, d-f), type II interferon receptor or interferon gamma receptor 1 (IFNγR1^-/-^, 4 to 18, g-i), or both (IFNαR1^-/-^ & IFNγR1^-/-^, n=5 or 6, j-l) were inoculated intravaginally with *C. trachomatis* serovar D at the dose of 2 x 10^5^ (a, d, g, & j), 5 x 10^6^ (b, e, h, & k) or 2 x 10^8^ (c, f, i, & l) inclusion forming units (IFUs) as indicated on top of the figure. All mice were monitored for live *C. trachomatis* recovery from the genital tract by taking vaginal swabs on days 1, 3, 5, 7, 10, 14, and weekly thereafter, as shown along the X-axis. The number of live *C. trachomatis* recovered from each swab at each time point was expressed as log_10_ IFUs, as displayed along the Y-axis. Data were acquired from three independent experiments. The data were compiled from 14 independent experiments. *p<0.05, Wilcoxon rank sum (2-tailed). IFUs from each time point or each group [in the form of area-under-curve (AUC)] were compared between the C57 control group and each gene-deficient mouse group. Note that IFNαR1^-/-^ mice significantly increased IFU shedding on days 3 and 5, but the shedding declined by day 7, while IFNγR1^-/-^ mice increased live chlamydial shedding on day 5. The increase persisted throughout the remainder of the infection course.

### 2. IFN**α**R1 also regulates *C. trachomatis* infection in the female genital tract at late stages

A careful comparison of shedding courses of live chlamydial organisms between IFNγR1^-/-^ mice and double knockout mice (deficient in both IFNγR1 & IFNαR1) revealed that although both IFNγR1^-/-^ and double KO mice significantly increased the yields of live *C. trachomatis* on days 5 or 7 postinfection, the shedding of live *C. trachomatis* from IFNγR1^-/-^ mice decreased after day 42 postinfection (Fig. 1). In contrast, the shedding of live *C. trachomatis* from the double KO mice remained high beyond day 42. The discrepancy was observed in mice receiving the *C. trachomatis* inoculum at either 5 × 10^6^ or 2 × 10^8^, suggesting that IFNαR1 signaling still plays a significant role in inhibiting *C. trachomatis* even at late stages of infection. This unexpected finding is consistent with the observation that double KO mice had a higher mortality rate than IFNγR1^-/-^ mice 6 weeks post-infection (Fig. 2). To further test the hypothesis, we evaluated whether IFNαR1 signaling can be blocked by an anti-IFNαR1 antibody (Fig. 3). A rat anti-IFNαR1 antibody treatment of C57 mice significantly increased the yield of live *C. trachomatis* in the vaginal swabs. In contrast, a normal rat IgG control failed to do so. The antibody-depletion results reproduced those obtained from IFNαR1^-/-^ mice (see Fig. 1, panel e vs. panel i), indicating that the anti-IFNαR1 antibody can be used to investigate the role of IFNαR1 signaling in chlamydial infection in the female genital tract. We then applied the same antibody treatment protocol to the IFNγR1^-/-^ mice 6 weeks after *C. trachomatis* infection (Fig. 4). The blockade of IFNαR1 signaling significantly increased the yield of live *C. trachomatis* from IFNγR1^-/-^ mice on days 54 and 56 post-infection when compared to that of IFNγR1^-/-^ mice treated with a normal rat IgG. Thus, we conclude that IFNαR1 signaling must also play a critical role in maintaining the inhibition of *C. trachomatis* in the female genital tract during the late stages of infection.

**Fig. 2.**
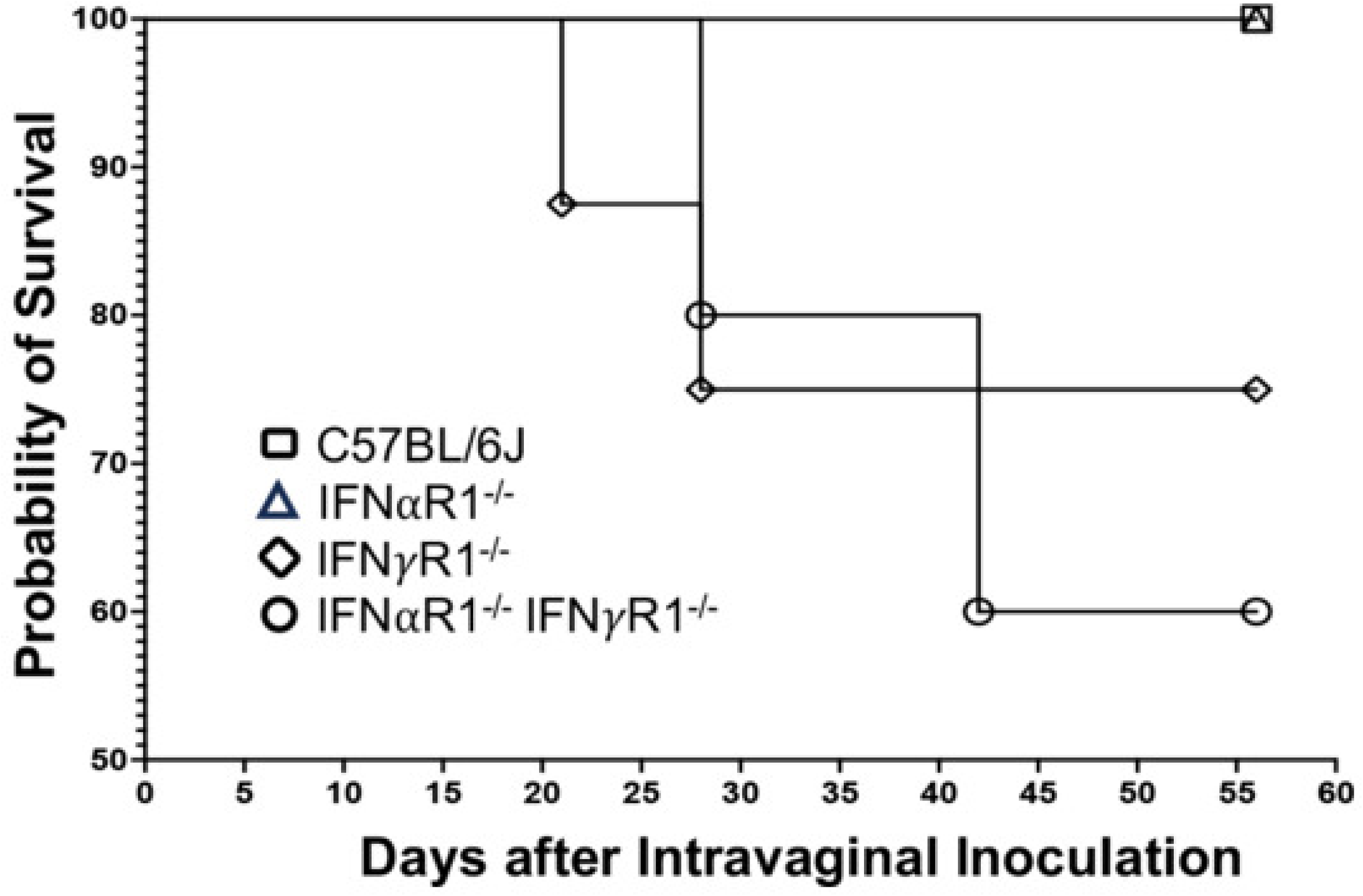
The effect of intravaginal infection with *C. trachomatis* on the survival probability of mice deficient in type I, type II, or both interferons. Groups of female mice without (C57BL/6J, n=22, open square) or with deficiency in type I interferon receptor (IFNαR1^-/-^, n=17, open triangle), type II interferon receptor (IFNγR1^-/-^, 18, open diamond), or both (IFNαR1^-/-^ & IFNγR1^-/-^, n=5, open circle) were inoculated intravaginally with *C. trachomatis* serovar D at the dose of 5 x 10^6^ IFUs as described in the legend of Fig.1. Mice were monitored for survival for 56 days as listed along the X-axis. The survival rate was displayed along the Y-axis. Note that mice deficient in IFNγR1 or both IFNαR1 & IFNγR1 died between 4 and 7 weeks post-infection.

**Fig. 3.**
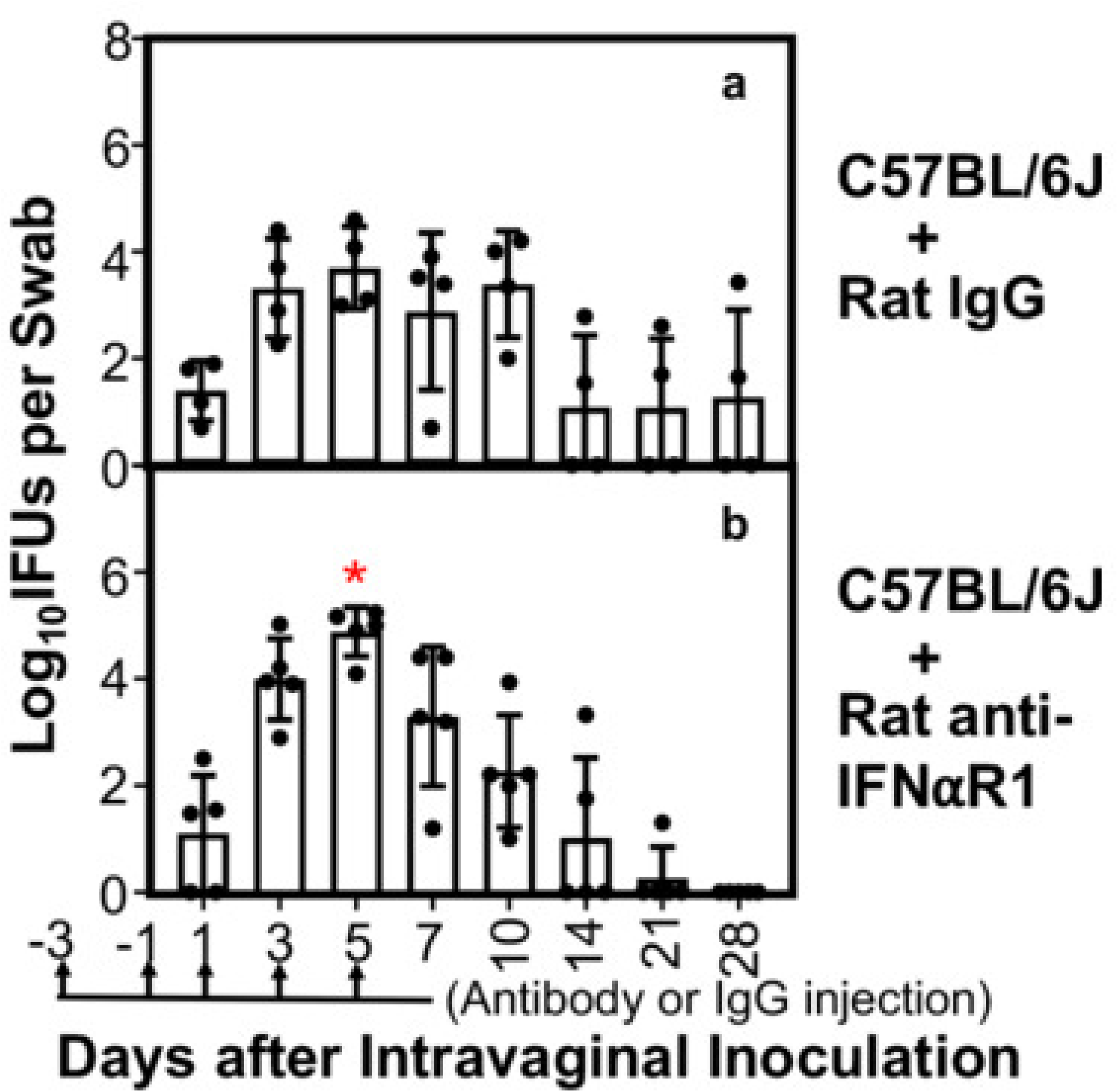
The effect of antibody blockade of IFNαR1 on *C. trachomatis* infection in the female genital tract of C57 mice. Female C57BL/6J mice treated with normal rat IgG (n=4, panel a) or rat anti-IFNαR1 antibody (n=5, b) were inoculated intravaginally with 5 x 10^6^ IFUs of *C. trachomatis* serovar D. All mice were monitored for live chlamydial organism shedding in vaginal swabs on days 1, 3, 5, 7, 10,14, 21 & 28, as listed along the X-axis. The number of live organisms recovered from each swab at each time point was expressed as log_10_ IFUs, and group means and standard deviation were displayed along the Y-axis. Data were acquired from two independent experiments. *p<0.05, Wilcoxon rank sum. IFUs from each time point were compared between the two groups. Note that the blockade of IFNαR1 increased the vaginal live chlamydial burden on day 5.

**Fig. 4.**
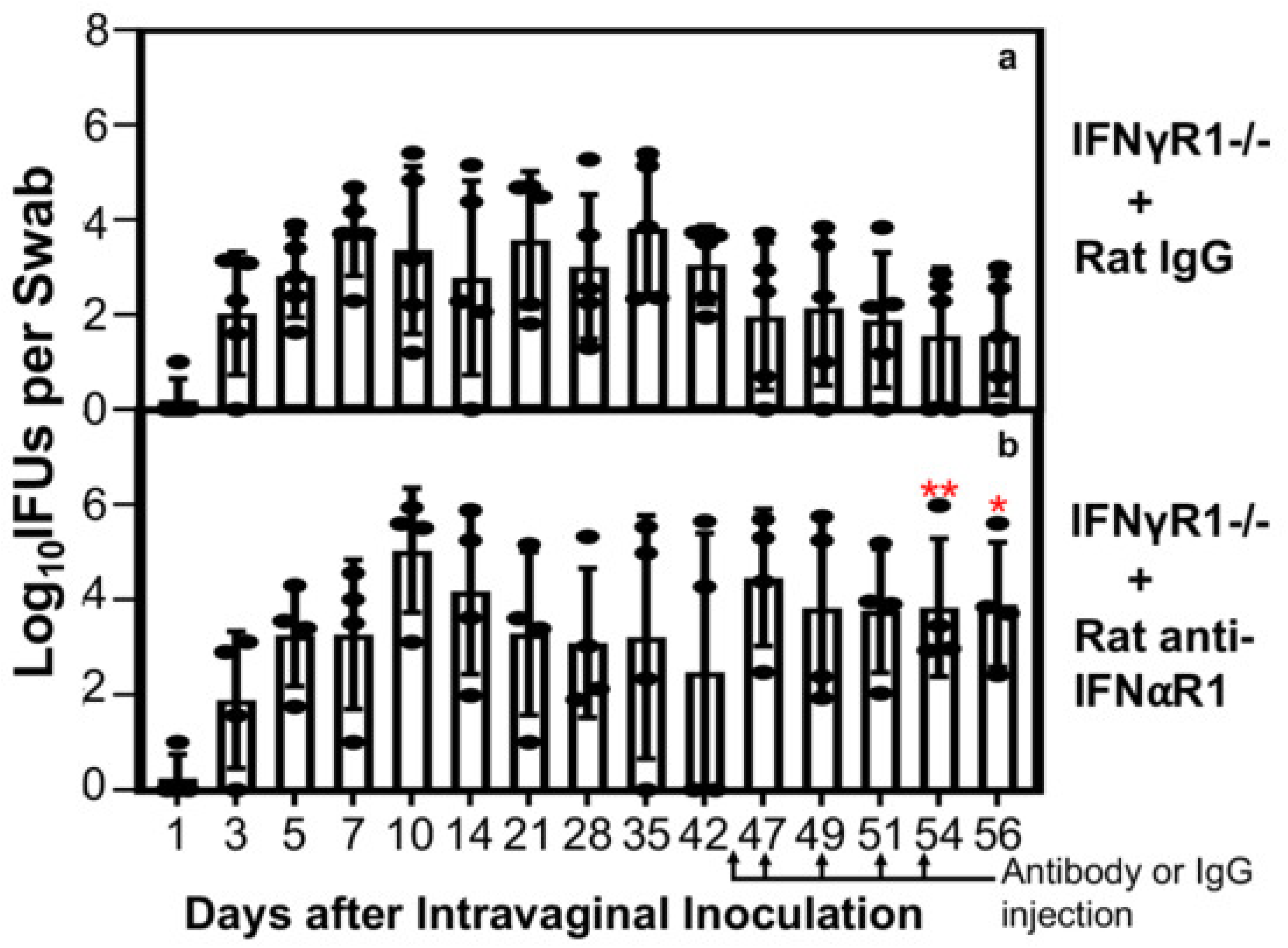
The effect of antibody blockade of IFNαR1 on *C. trachomatis* infection in the female genital tract of IFNγR1^-/-^ mice. nine female IFNγR1^-/-^ mice inoculated intravaginally with 5 x 10^6^ IFUs of *C. trachomatis* serovar D were monitored for live organism shedding in vaginal swabs on days 1, 3, 5, 7, 10,14, and weekly thereafter, and as listed along the X-axis. On day 42, the mice were randomly divided into two groups for treatment with rat IgG (n=5, panel a) or rat anti-IFNαR1 antibody (n=4, panel b) on days 45, 47, 49, 51 & 54, respectively, as indicated along the X-axis. Additional vaginal swabbing was carried out on days 47, 51 & 54 beyond the original weekly schedule to more accurately evaluate the effect of antibody treatment on *C. trachomatis* shedding. The number of live organisms recovered from each swab at each time point was expressed as log_10_IFUs, and group mean and standard deviation were displayed along the Y-axis. Data were acquired from two independent experiments. *p<0.05 while **p<0.01, Wilcoxon rank sum. IFUs from each time point were compared between the two groups. Note that the blockade of IFNαR1 increased the live chlamydial burden on days 54 & 56.

### 3. The IFN**α**R regulates *C. trachomatis* in the lower vagina, while IFN**γ**R does it in the upper vagina

Having demonstrated both distinct and overlapping roles of IFNαR1 signaling and IFNγR1 signaling in controlling *C. trachomatis* infection in the female genital tract over the infection time course, we next tested whether IFNαR1- and IFNγR1-dependent inhibition of *C. trachomatis* is restricted to different segments of the female genital tract. As shown in Fig. 5, compared to the wild-type C57 mice, IFNαR1^-/-^ mice significantly increased the yield of live *C. trachomatis* in the lower vaginal tissue, while IFNγR1^-/-^ mice significantly increased the yield of live *C. trachomatis* in the upper vaginal tissue (left panels) on day 5 after the intravaginal inoculation. This observation was validated by simultaneously monitoring chlamydial genome copies (right panels). Significantly elevated levels of chlamydial genomes were recovered from the upper vaginal and ecto-cervical tissues of the IFNγR1-/- mice, while the IFNαR1-/- mice significantly increased the chlamydial genocopies in the lower vaginal tissue only. Neither live chlamydial organisms nor chlamydial genomes detected in other genital tissues differed significantly across mouse groups. These observations together suggest that IFNαR1 signaling inhibits *C. trachomatis* in the lower vagina, while the inhibition of *C. trachomatis* by IFNγR1 signaling is restricted to the upper vagina-ectocervix regions.

**Fig. 5.**
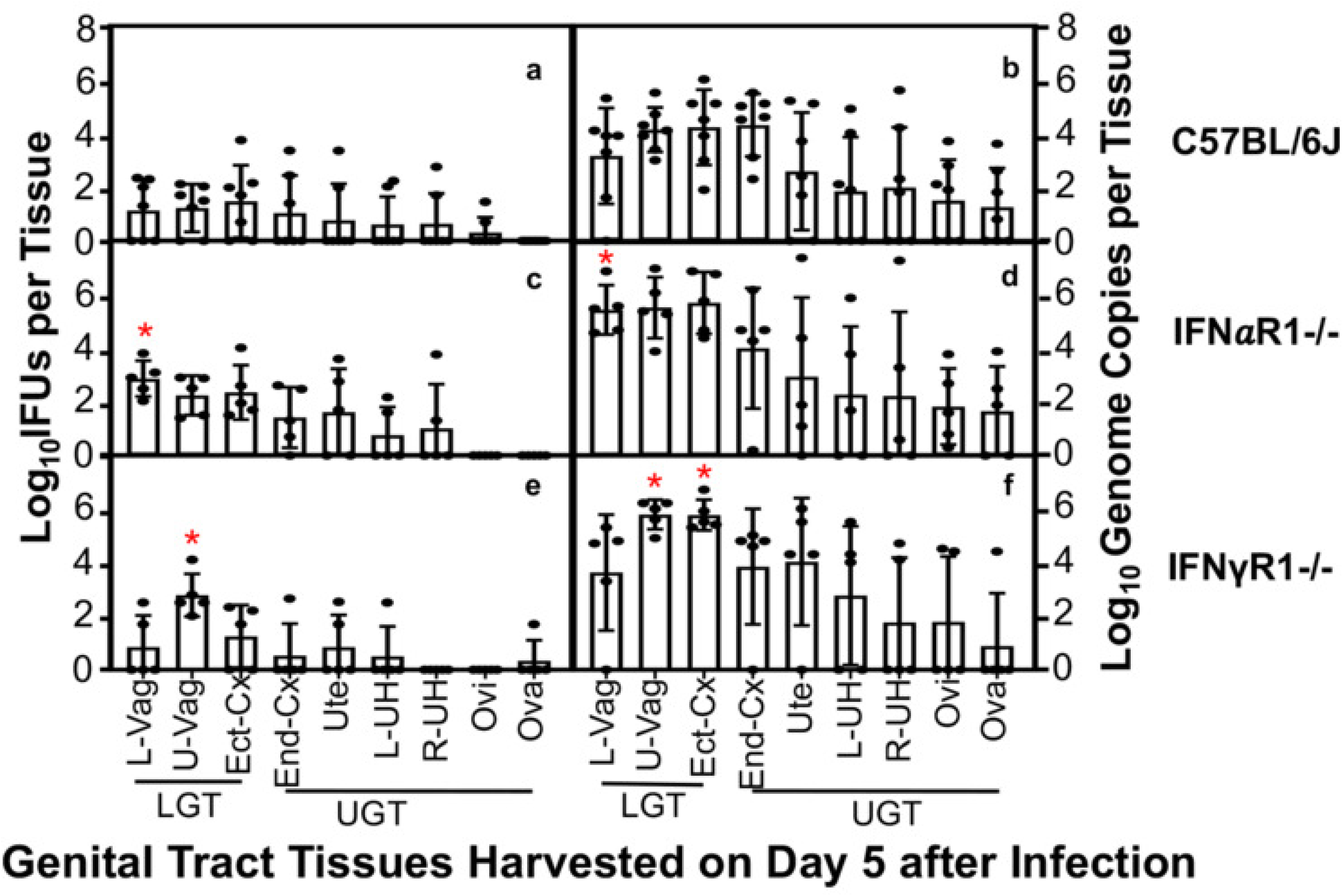
Chlamydial burdens from different genital tract tissues of mice with or without deficiency in type-I or type-II interferons. Female mice without (C57BL/6J, n=7, panels a & b) or with deficiency in type-I interferon receptor (IFNαR1^-/-^, n=5, c & d), type-II interferon receptor (IFNγR1^-/-^, n=5, e & f) were inoculated intravaginally with *C. trachomatis* serovar D at the dose of 5 x 10^6^ IFUs. All mice were euthanized on day 5 after inoculation to quantify chlamydial organisms from different segments of both the lower genital tract (LGT) and upper genital tract (UGT), as listed along the X-axis (see the Materials and Methods section for specific tissue segment). The chlamydial burdens were quantified as live organisms (IFUs, a, c, & e) and total organisms (genome copies, b, d, & f) and shown along the left and right Y-Axes, respectively. Data were acquired from three independent experiments. *p<0.05, Wilcoxon rank sum. IFUs or genome copies from each genital tissue were compared between C57 and each gene-deficient group. Note that significantly more chlamydial organisms were recovered from the lower vagina of IFNαR1^-/-^ and upper vagina/ectocervix of IFNγR1^-/-^ mice.

## Discussions

Chlamydial infection induces both type I (17–19) and type II (34, 40, 43–48) IFNs, and their roles in chlamydial infection have been demonstrated in various models (21–24, 34, 43, 45–47, 49). This study was designed to elucidate the relationships between the two IFN systems with respect to their anti-chlamydial activities in the female genital tract across time and space at a wide range of inoculation doses. First, along the infection time course, we found that signaling by type I interferon receptor IFNαR1, but not the type II interferon receptor IFNγR1, played a dominant role in inhibiting *C. trachomatis* infection during the early stage. IFNαR1^-/-^ mice significantly increased the yield of live chlamydia on days 3 & 5 post-inoculation, while the earliest increase in chlamydial yield from IFNγR1^-/-^ mice was detected on day 5 under multiple inoculation doses. Further, the anti-chlamydial activity of IFNαR1 signaling was transient, whereas that of IFNγR1 signaling was long-lasting, as IFNαR1^-/-^ mice reduced their shedding level to that of wild-type mice by day 7, while IFNγR1^-/-^ mice maintained a significant increase in the chlamydial yield for the remainder of the time course. These observations together demonstrate a temporal division of labor between type I & II interferon-mediated anti-chlamydial activities. Second, we surprisingly found that IFNαR1 signaling is still able to inhibit *C. trachomatis* at late stages of infection, as the double-knockout mice (deficient in both IFNαR1 & IFNγR1) developed a higher rate of mortality and shed more live chlamydial organisms than IFNγR1^-/-^ mice 6 weeks after inoculation. This conclusion was validated by blocking IFNαR1 signaling with an anti-IFNαR1 antibody, which significantly increased genital shedding of live *C. trachomatis* in IFNγR1^-/-^ mice. Third, the anti-chlamydial activity of IFNαR1 signaling is localized to the lower vagina. In contrast, the anti-chlamydial activity of IFNγR1 signaling is localized to the upper vagina. These observations demonstrate a spatial division of labor between type I & II interferon-mediated anti-chlamydial activities. Thus, both type I & II interferon signaling systems are important in controlling *C. trachomatis* infection in the female genital tract by complementing and compensating for each other to maintain sustained pressure on the infection.

The dominant role of IFNαR1 signaling but not of IFNγR1 signaling on day 3 after intravaginal inoculation with *C. trachomatis* may reflect the rapid induction of type I IFN over type II IFN at the early stage of infection. Chlamydia can activate cGAS-STING (50), TLR3 (51–53), or TLR2 (54) to induce type I IFN production. Mice deficient in either cGAS or STING were more susceptible to *C. trachomatis* infection, with a significantly increased yield of live *C. trachomatis* in vaginal swabs on days 3 and 5 post-infection (20). Consistently, IFNαR1^-/-^ mice also increased the yield of live *C. trachomatis* during the early stage of infection, as reported previously (21) and in the current study. In contrast, mice deficient in type II IFN receptor (IFNγR1^-/-^) delayed the increase in the yield of live *C. trachomatis* to day 5 postinfection. This may partially reflect a slower induction of IFNγ following an intravaginal infection with *C. trachomatis*. Although trans-cervical inoculation with *C. trachomatis* has been shown to induce ILC3s to produce IFNγ as early as day 3 post-inoculation (32), it represents a more severe stress than intravaginal inoculation, thereby accelerating the induction of innate IFNγ. This is consistent with a previous finding that transcervical inoculation consistently induced IFNγ-dominant responses (55). More importantly, the rapid early production of type I IFNs may suppress the expression and anti-chlamydial function of IFNγ, as IFNα/β has been shown to inhibit IFNγ expression by NK and T cells (56). This inhibition depends on IFNαR1 signaling and STAT1. Thus, the temporal division of labor between type I and II interferons in blocking chlamydial infection reflects both the genital tract’s responsiveness to intravaginal inoculation with *C. trachomatis* and the interactions between these two interferon systems. As a result, the decline in the production of type I IFNs will ensure a timely replacement of the IFNαR-dependent anti-chlamydial activity by the antichlamydial activity mediated by IFNγR signaling. Intravaginal inoculation with *C. trachomatis* may impose a stress sufficient to induce type I IFNs via mechanisms such as the cGAS/STING pathway, leading to the inhibition of both *C. trachomatis* and IFNγ expression. The decline in anti-chlamydial activity mediated by IFNαR1 signaling may lead to the removal of the type I IFNs’ inhibitory effect on IFNγ expression by cells such as NK cells. Thus, the IFNαR1’s anti-chlamydial activity is replaced by that of innate IFNγ first, followed by IFNγ produced by *C. trachomatis* epitope-specific lymphocytes. The switch in anti-chlamydial activity between type I and II is also consistent with the concept that an appropriate host response may both maintain the anti-microbial activity and minimize tissue damage. Although the precise mechanism of IFNαR1 signaling’s anti-chlamydial activity remains unknown, IFNγR1 signaling is effective in controlling chlamydial infection (xx), which can be maintained by *C. trachomatis* epitope-specific lymphocytes. Thus, a sustained anti-chlamydial activity via IFNγR1 signaling is more beneficial for the host.

The finding that IFNαR1 signaling can still inhibit *C. trachomatis* at late stages of infection is somewhat unexpected, as type I IFNs typically appear early during microbial infection and inflammation. We made this finding by carefully comparing mortality rates and the recovery of live *C. trachomatis* from vaginal swabs between IFNγR1^-/-^ and double-knockout mice over 8 weeks. The double-knockout mice exhibited a higher mortality rate and higher levels of live *C. trachomatis* shedding by week 6, which coincided with a significant decline in live *C. trachomatis* shedding in IFNγR1^-/-^ mice. The reduction in live *C. trachomatis* shedding 6 weeks after infection may suggest that IFNγR1^-/-^ mice may have developed IFNγ-independent anti-chlamydial mechanisms by this time. Type I IFN may represent a major contributor to the IFNγ-independent anti-chlamydial mechanisms, as the double-knockout mice still maintained high levels of live *C. trachomatis* shedding. This conclusion was validated by the observation that antibody blockade of IFNαR1 signaling significantly increased live *C. trachomatis* shedding in IFNγR1^-/-^ mice. Thus, IFNαR1 signaling can still inhibit *C. trachomatis* replication at late stages of infection in the female genital tract. Clearly, more experiments are required to fully reveal the significance of IFNαR1 signaling-mediated anti-chlamydial activity during the late stages of infection.

The current study has also, for the 1^st^ time, demonstrated a clear spatial division of labor between type I and II interferon-mediated anti-chlamydial activities. The anti-chlamydial activity of IFNαR1 signaling is localized to the lower while that of IFNγR1 signaling to the upper vaginal tissues on day 5 postinfection, when the anti-chlamydial activities of both receptors are measurable by monitoring the yield of live *C. trachomatis* in the vaginal swabs. This finding is consistent with our previous reports that mice deficient in either the production of type I IFNs (20) or the responsiveness to type I IFNs (21) allowed more *C. trachomatis* replication in the lower, but not upper, genital tract of the female mice. The localization of the type I IFN-dependent anti-chlamydial activity to the lower genital tract was not caused by the intravaginal inoculation, as transcervically inoculated mice also displayed type I IFN-dependent anti-chlamydial activity in the lower genital tract. The current study more precisely mapped type I IFN-dependent anti-chlamydial activity to the lower vagina. This information may help guide the design of vaginal strategies for reducing the transmission of sexually transmitted infections by enhancing type I IFNs in the lower genital tract. More significantly, type II IFN-dependent anti-chlamydial activity was localized to the upper vagina and ectocervix on day 5 following an intravaginal inoculation. This finding suggests that the upper vagina and ectocervix of the female genital tract may be targeted for priming chlamydial epitope-specific IFNγ production via vaccination to prevent ascending infection and upper genital complications.

## Materials and Methods

### 1. Mouse infection with *Chlamydia trachomatis*

*Chlamydia trachomatis* serovar D organisms (clone# UW-3/Cx, ATCC# VR-885) were acquired from ATCC (Manassas, VA 20110). The chlamydial organisms were amplified/passaged in HeLa cells (human cervical carcinoma epithelial cells; ATCC# CCL-2), and purified into elementary bodies (EBs) using discontinuous density centrifugation (57). The purified EBs were kept in aliquots at -80°C until use.

The mouse experiments were carried out in accordance with the recommendations in the Guide for the Care and Use of Laboratory Animals endorsed by the National Institutes of Health (58). The protocol was approved by the Committee on the Ethics of Laboratory Animal Experiments of the University of Texas Health Science Center at San Antonio. Mice used in the current study were female, 7-10 weeks old, and purchased from Jackson Laboratories, Inc. (Bar Harbor, ME). They include wild-type mice (C57BL/6J, stock# 000664), or mice deficient in type I interferon receptor IFNαR1 [IFNαR1^-/-^ or B6(Cg)-*Ifnar1^tm1.2Ees^*/J, stock# 0285288], or type II interferon receptor IFNγR1 (IFNγR1-/-, B6.129S7-*Ifngr1^tm1Agt^*/J, stock# 003288), or both (double-knockout, IFNαR1-/- & IFNγR1-/-, B6.Cg-*Ifngr1^tm 1Agt^ Ifnar1^tm1. 2 Ees^* /J, stock# 029098). Mice were inoculated intravaginally with *C. trachomatis* EBs at 2 x 10^5^, 5 × 10^6^, or 2 x 10^8^ IFUs (inclusion-forming units) per mouse as described previously (59). Briefly, EBs were diluted in 10 µl of sucrose-phosphate-glutamic acid (SPG) buffer (consisting of 220 mM sucrose, 12.5 mM phosphate, 4 mM L-glutamic acid, pH 7.5) and delivered gently to the ectocervix using a p20 pipettor tip. After inoculation, mice were monitored for live chlamydial shedding and/or chlamydial genome copies in vaginal swabs or sacrificed to titrate live organisms and genomes in designated organs/tissues.

### 2. Collection of vaginal swabs and genital tract tissue samples from mice

To monitor live chlamydial shedding from the female genital tract, vaginal swabs were taken on days 1, 3, 5, 7 & 10 and weekly thereafter after infection. Each swab was collected into an Eppendorf vial containing 0.5 ml of SPG buffer and 3-5 glass beads. The vials were kept on ice until vaginal swabs from all mice were collected. Each swab-containing vial was thoroughly vortexed to release infectious EBs into the supernatants. To titrate live chlamydial organisms recovered from different segments of the female mouse genital tract tissues, parallel groups of mice were sacrificed on day 5 after the intravaginal inoculation and the following genital segments were collected: The lower genital tissues (LGT) include lower vagina (L-Vag), upper vagina (U-Vag), and ectocervix (Ect-Cx), while the upper genital tissues (UGT) include endocervix (End-Cx), uterus (Ute), left uterine (L-UH), right uterine (R-UH), oviduct (Ovi), and ovary (Ova). Each tissue segment was transferred to an Eppendorf vial containing 0.5 ml cold SPG buffer for tissue homogenization and sonication.

### 3. Infect HeLa cell monolayers grown in 96-well microplates with mouse samples to quantify live *C. trachomatis*

The vortexed swab-containing vials and the sonicated tissue-containing vials were briefly centrifuged to pellet cell debris. The live chlamydial organisms in the supernatants were titrated on HeLa cell monolayers grown in 96-well microplates in duplicate as described previously (60). Briefly, each swab sample was diluted into 1:0 (neat), 1:4, 1:16, 1:64, and 1:256 using SPG in a 96-well dilution microplate. Then, 100 μL from each dilution was transferred to the HeLa monolayer plate. It is important that the HeLa monolayers be less than 24 hours old and at ∼70% confluence before use, which should improve sensitivity and reproducibility. To further improve sensitivity, the HeLa monolayers were pretreated with DEAE for 10 min prior to inoculation with chlamydial samples, and the inoculated microplates were centrifuged to increase the contact between chlamydial organisms and HeLa cells. The infected HeLa cells were incubated at 37 °C for 48 h in a CO2 incubator before processing for immunofluorescence assays.

### 4. Immunofluorescence assay

The infected HeLa cell monolayers were processed for immunofluorescence detection of *C. trachomatis* as described previously (61). Briefly, HeLa cells grown on coverslips were fixed with paraformaldehyde (Sigma) and permeabilized with saponin (Sigma). After washing and blocking, the cell samples were subjected to a combination of antibody and chemical staining. Hoechst (blue; Sigma) was used to visualize nuclear DNA. A rabbit anti-chlamydial antibody (raised by immunization with serovar D EBs [data not shown]) plus a goat anti-rabbit IgG conjugated with Cy2 (green; Jackson ImmunoResearch Laboratories, Inc) were used to visualize chlamydial inclusions. The immunolabeled monolayers were used to count chlamydial inclusions under an Olympus IX-81 fluorescence microscope equipped with multiple filter sets (Olympus, Melville, NY).

### 5. Calculation of live C. trachomatis titers from each vaginal swab or tissue segment

The green fluorescence-labeled chlamydial inclusions were counted under the appropriate objective lens. Randomly selected 5 views from each well were counted to determine the total number of inclusions per well. When the number of inclusions per well was few enough, the entire well was counted. The number of inclusions per well was used to calculate the number of inclusions per swab or tissue segment sample, accounting for factors such as sample dilution and the amount of each sample used to inoculate HeLa cells. The total number of IFUs/swab or tissue was converted to a log_10_ value to calculate the group mean and standard deviation at each time point. The detection limits of the above titration method were 5 IFUs per swab or tissue sample. This is because 100 µl from the total volume of 500 µl per neat sample or each serially diluted sample was used to inoculate HeLa cell monolayers.

### 6. Quantitative PCR for measuring chlamydial genomes in each genital tissue segment

After a genital tissue segment was homogenized in 500 μL of SPG in an Eppendorf vial, 100 μL of the tissue lysate was transferred to a fresh Eppendorf vial for DNA extraction using the QIAamp DNA Mini Kit-250 (Qiagen, Frederick, MD) according to the manufacturer’s instructions. Each DNA preparation was collected into 100 μl of water, and 4 μl of each DNA sample was used for quantitative PCRs (qPCRs) using the following primers complementary to chlamydial 16S rRNA coding region: The forward primer is 5′-CGCCTGAGGAGTACACTCGC-3′, and a reverse primer is 5′-CCAACACCTCACGGCACGAG-3′, for amplifying a 208-bp fragment. The probe primer is 5′-CACAAGCAGTGGAGCATGTGGTTTAA-3′. All primers were synthesized by Integrated DNA Technologies (Coralville, IA). A plasmid containing the 208-bp fragment of the chlamydial 16S ribosomal gene was used as a standard template for quantification. PCR was performed in a total volume of 20 μl using a CFX96 Touch Deep Well Real-Time PCR Detection System with iQ Supermix real-time PCR reagent (Bio-Rad, Hercules, CA). The PCR conditions were as follows: initial denaturation at 95°C for 3 min, followed by 40 cycles of amplification at 95°C for 15 s and 60°C for 1 min. The results were expressed as the total number of genome copies per sample and plotted on a log_10_ scale.

### 7. Statistics

The numbers of live chlamydial organisms (expressed as log_10_ IFUs) or log_10_ genome copies per sample/mouse, were compared between wild-type mice and mice deficient in type I and/or II interferon receptors using the Wilcoxon rank-sum test (2-tailed). An ANOVA was first performed to assess the overall difference among the three groups before comparing any two groups. Wilcoxon rank sum test is used to compare both nonparametric and parametric data (62).

## ACKNOWLEDGMENT

This work was supported in part by grants from the U.S. National Institutes of Health (U01AI182210 to G.Z.).

